# Fungal Ice2p is in the same superfamily as SERINCs, restriction factors for HIV and other viruses

**DOI:** 10.1101/2021.01.08.425964

**Authors:** Ganiyu O. Alli-Balogun, Tim P. Levine

**Author notes:** **Data availability statement:** Data that support this study are freely available in Harvard Dataverse at https://dataverse.harvard.edu/dataverse/Ice2_SERINC.

## Abstract

Ice2p is an integral endoplasmic reticulum (ER) membrane protein in budding yeast *S. cerevisiae* named ICE because it is required for Inheritance of Cortical ER. Ice2p has also been reported to be involved in an ER metabolic branch-point that regulates the flux of lipid either to be stored in lipid droplets or to be used as membrane components. Alternately, Ice2p has been proposed to act as a tether that physically bridges the ER at contact sites with both lipid droplets and the plasma membrane via a long loop on the protein’s cytoplasmic face that contains multiple predicted amphipathic helices. Here we carried out a bioinformatic analysis to increase understanding of Ice2p. Firstly, regarding topology, we found that diverse members of the fungal Ice2 family have ten transmembrane helices, which places the long loop on the exofacial face of Ice2p, where it cannot form inter-organelle bridges. Secondly, we identified Ice2 as a full-length homologue of SERINC (serine incorporator), a family of proteins with ten transmembrane helices found universally in eukaryotes. Since SERINCs are potent restriction factors for HIV and other viruses, study of Ice2p may reveal functions or mechanisms that shed light on viral restriction by SERINCs.

## Introduction

Many functions of membrane proteins remain far more mysterious than their soluble counterparts largely because the hydrophobic environment remains hard to probe. One way to make progress in understanding membrane proteins has been to determine their structures, with a step-change achieved through single particle cryo-electron microscopy ^1,2^. Another approach is to apply bioinformatic techniques to a protein of interest for which information is lacking to link it to others that are better undestood.^3^ Likely discoveries here include finding topological features that indicate a limited range of functions, identifying homologies where sequence has diverged below the threshold required for conventional tools, and defining the arrangement of transmembrane helices (TMHs) from *ab initio* modelling based on pairwise evolution of contacting side-chains.^4^

Ice2p is a multi-spanning ER localised yeast protein of 491 residues that was first named for its role in <u>i</u>nheritance of <u>c</u>ortical <u>E</u>R from mother to daughter cells.^5^ Cortical ER is reduced particularly in buds upon deletion of *ICE2* alone,^5^ and it is markedly reduced in both mother and daughter cells by deleting *ICE2* in strains already carrying other mutations that reduce cortical ER.^6,7^ The molecular function of Ice2p is unknown and genetic interactions implicate a wide range of ER-related activities. These include an indirect effect on protein targeting to the inner nuclear envelope,^8,9^ a role in stabilising ER membrane proteins and ER-associated degradation,^10–12^ and supporting the function of some ER membrane proteins.^13^

While none of these links are direct, multiple strands of evidence link Ice2p to lipid metabolism, with some direct interactions. Ice2p is required for the mobilisation of triacylglycerol in lipid droplets by conversion first to diacylglycerol then phospholipids as bilayer components during the resumption of growth after stationary phase.^14^ Thus, *ice2*∆ cells are slower to re-enter log-phase growth. Conversion of neutral lipid to phospholipid may underlie the requirement for Ice2p in the induction of macroautophagy.^15^ A more recent study focussed on Ice2p because of its role in supporting ER expansion, and showed a direct interacts between Ice2p and Spo7p (the yeast homologue of human TMEM18A, also called NEP1-R1), which inhibits Spo7p and thereby indirectly inhibits Pah1p (homologue of lipin in humans).^16^ Since Pah1p/lipin channels phosphatidic acid, the archetypal membrane-building phospholipid, into diacylglycerol for storage in lipid droplets,^17^ Ice2p functions to promote membrane biogenesis in the ER.

Alongside a biochemical role in lipid metabolism, Ice2 has a physical relationship with lipid droplets. In cells in stationary phase GFP-tagged Ice2p localises to parts of the ER that contact lipid droplets, and it dissociates rapidly into general ER after re-initiation of growth.^14^ Lipid droplet localisation was attributed to direct bridging by a long loop predicted by the topology engine TMHMM2.0 to be in the cytoplasm between TMH6 and TMH7 among eight TMHs overall, revising an original designation of seven TMHs.^5^ The long loop on its own was sufficient for lipid droplet targeting both in yeast and in COS7 cells, and within that loop four regions were identified as forming amphipathic helices, making them candidates to bind the lipid droplet surface.^14,18,19^ The proposed bridging from ER to lipid droplets was subsequently extended to suggest that Ice2 also direct bridges from ER to plasma membrane to form cortical ER.^7^

Given that the molecular function of Ice2p is unknown, a bioinformatics approach might reveal useful insights. Here we first analysed the topology of Ice2 in context of the whole Ice2 family, revising the number of TMHs to ten, which places the long loop in the ER lumen, rendering it unable to form bridges, so largely ruling out that Ice2p tethers the ER either to lipid droplets or to the plasma membrane. Secondly, we found that Ice2p is a full-length homologue of SERINCs, a ubiquitous eukaryotic family of proteins also with ten TMHs, named for their possible role in <u>ser</u>ine <u>inc</u>orporation into lipids, and best known for their role as potent restriction factors for HIV and other viruses.^20,21^ Given the conserved elements identified in a cryo-electron microscopy structure of SERINC,^22^ this allowed us to identify likely key functional residues in Ice2 that can inform future experiments.

## Methods

### Alignment of yeast Ice2 sequences

An alignment of 18 Ice2 homologues was made from experimentally relevant and diverse Ice2-positive fungi (Supplementary Figure 1). Sequences were aligned by Clustal Omega, and a tree created with PHYML 3.0.^23,24^ A medium-sized group of 95 Ice2 sequences was obtained from UNIPROT by filtering (i) to remove fragments; (ii) to include only sequences between 200 and 800 residues; (iii) using UniRef clusters with maximum identify 90% (nr90), so that no sequence shares >90% identity to another.

### Amphipathic helix prediction

Sequences of amphipathic helices previously identified in Ice2p were plotted as cartwheels using the Heliquest server.^14,25^ Heliquest was also used to calculate the hydrophobicity and hydrophobic moment of each region, as well as additional hydrophobic regions in Ice2p.

### Topology predictions

Transmembrane helix prediction tools were used via online servers: TOPCONS, TMHMM2.0 and MemBrain using default parameters.^26–28^ For TOPCONS, reports include the consensus and the five component programmes: OCTOPUS, Philius, Polyphobius, SCAMPI, and SPOCTOPUS.

### Iterative searching with Jackhmmer

Profiles based on hidden Markov models were built by iterative searches by Jackhmmer, part of the HMMER suite, using standard settings, E-values = 0.01 for the whole sequence, 0.03 for each hit.^29^

### Profile-profile searches and structure prediction

HHpred, an online enactment of HHsearch at the Tuebingen toolkit was used for profile-profile searches for Ice2p homologues in PFAM, in PDB and in model eukaryotes including human and yeast.^23,30,31^ Settings were standard, except initially we used eight iterations of HHblits, and e-value for inclusion was 0.01. Searches in FFAS, used standard settings.^32^

For modelling a structure for Ice2p based on the structure of *D. melanogaster* SERINC, the HHpred alignment was extended beyond the regions of highest homology by setting alignment mode to local realignment and setting the MAC realignment threshold to 0.01. This alignment was forwarded to Modeller ^33^. Other template-based tools used to build 3D models of Ice2 were Phyre2 (intensive mode), Galaxy and SWISS-MODEL(standard settings).^34–36^

Models made by analysis of contact co-evolution were made in trRosetta, switched either to ignore known structures or to use them as templates.^37^ For making models with full control of the multiple sequence alignment (MSA), RaptorX was initiated with a pre-aligned list of 1202 Ice2 homologues uncontaminated with any SERINCs.^38^ This list was obtained by submitting the 61 hits of the 3^rd^ round of HHblits with Ice2 (351 aa, missing inserts, Supplementary Table 2D) to a single round of PSI-BLAST in the NCBI nr100 database, with E-value ≤1×10^−5^ to prevent inclusion of any SERINCs, and then deleting columns so that Ice2 was ungapped.

### Phylogeny Tree and Cluster analysis

A list of 223 sequences (Supplementary Table 4) was accumulated from three sources: (i) 184 sequences including diverse SERINCs: prepared by seeding the NCBI PSI-BLAST server with *Thecomonas trahens* hypothetical protein AMSG_07160 (XP_013756618.1; 338 aa), restricting results to 29 taxonomic groups, otherwise with standard settings (*e.g.* threshold p=0.005). Convergence was obtained at iteration 9, the included 469 sequences were aligned sequences by Clustal Omega, the alignment was edited by hand to remove long, unique insertions, fragments and highly similar repeats, leaving 184 sequences. These included 167 SERINCs, 9 fungal Ice2s, and 8 others with no known domain, including the seed. (ii) 34 further sequences, documented as either Ice2 or with no known domain, obtained from the second round of PSI-BLAST set without limit on the taxonomy of hits (unlike (i) above) seeded either with the *T. trahens* protein as (i) or with the *Nematostella vectensis* predicted protein EDO39878.1 (XP_001631941.1, 395 aa). (iii) Five further sequences obtained by hand curating incomplete sequences in (i) and (ii)..

For a phylogeny tree, sequences were aligned with Kalign, and then relationships were identified by PHYML3.0 with branch support by aBayes.^24,39^. To identify relationships in an all-*vs.*-all cluster map, sequences were submitted CLANS with standard settings (threshold e-values: 1e^−4^).^40^

## Results

### The Ice2p region identified as a loop may contain at least one TMH

*S. cerevisiae* Ice2p was initially predicted to span the ER membrane with 7 TMHs,^5^ and later this was revised to 8 TMHs using TMHMM (Figure 1A).^14,27^ To determine the most likely topology of Ice2p, we considered that the topology was highly likely to be conserved across the whole family of proteins related to Ice2, known as protein family PF08426 with single proteins per fungal species in all dikarya (Ascomycota and Basidiomycota) but absent from most Mucormycota and all Chytridiomycota, Zoopagomycota and Microsporidia (not shown). 41 Importantly, homology is present across the full length of diverse Ice2 family members, and the most obvious outlier is *S. cerevisiae* Ice2p because of two non-conserved insertions: 106-137 and 292-343 (Supplementary Figure 1A/B). The second insertion is entirely within the region 241-357 previously described as a cytoplasmic loop that contacts and binds to lipid droplets (Figure 1B).^14^ The loop is predicted to be largely helical (Figure 1A) and was proposed to contain four predicted amphipathic helices (AH1-4), using Heliquest (Figure 1B).^25^

**Figure 1.**
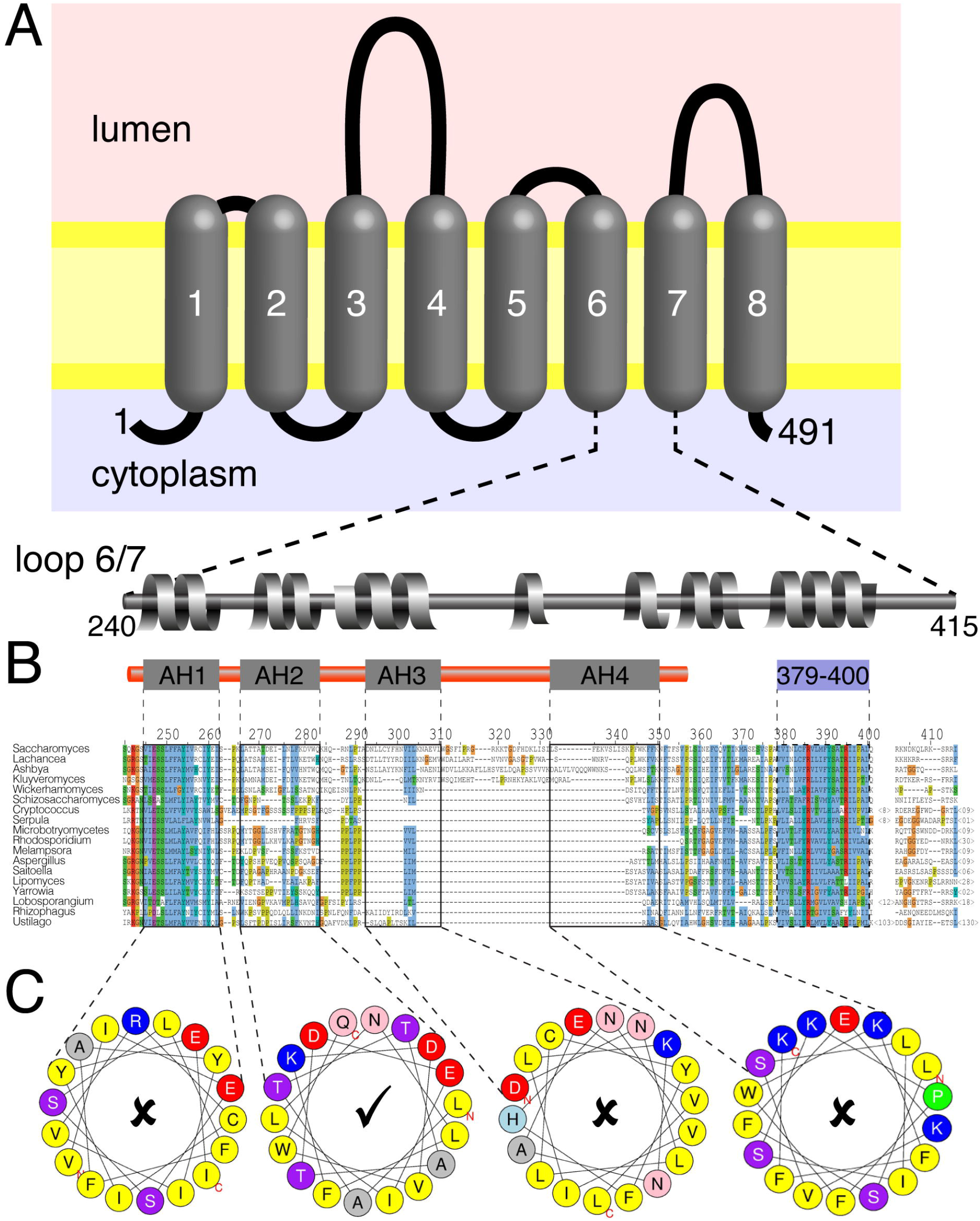
Analysis of loop between previously predicted TMH6 and 7 of Ice2. A. Topology of Ice2p predicted by TMHMM2.0: eight transmembrane helices (TMH) with residues 240-415 forming a cytoplasmic loop between TMH6 and TMH7.^14^ Also shown are α-helical regions in that loop predicted by PSIPRED3.0. B. Alignment of residues 240-414 in Ice2p with homologues from 17 other fungal species (see also Supplementary Figure 1). Residues 241-357 indicated by red bar (top), previously shown to target lipid droplets (Markgraf et al., 2014), contain four regions previously described as amphipathic helices AH1-4 (boxes). In addition, residues 379-400 are indicated by blue bar (top) and dashed box. Alignment coloured according to CLUSTALX. C. Helical wheel representations generated in Heliquest for AH1-4. Tick (✓) on helical wheel indicates suitability to fold into an AH, while crosses (x) indicate hinderance to AH formation because of physico-chemical parameters more characteristic of a TMH (high hydrophobicity, low hydrophobic moment, low incidence of non-polar residues, see Supplementary Table 1), or presence of proline, or a polar residue within the predicted hydrophobic interface.

We re-examined the predictions for AH1-4, in particular to distinguish between an AH and a TMH from biophysical characteristics returned by Heliquest: total hydrophobicity (TMH higher than AH), hydrophobic moment (TMH lower than AH) and proportion of polar residues (TMH lower than AH). While AH2 (residues 266-283) has all of the parameters characteristic of an AH, all three parameters for AH1 (residues 245-262) are characteristic of a TMH (Supplementary Table 1, Figure 1C).^42,43^ Furthermore, AH3 (residues 293-310) has none of the AH parameters, and AH4 (residues 333-350) has intermediate parameters, but it contains a helix breaking proline (Pro344), and the middle of the proposed hydrophobic face contains a serine residue (S342), making it unlikely to form an AH (Supplementary Table 1, Figure 1C). Both AH3 and AH4 are in the portion of Ice2p without counterparts in most fungi (Figure 1B), which indicates that any fundamental conserved function of the Ice2 family is unlikely to involve these regions. This analysis indicates that the region of Ice2p expressed by Markgraf *et al*. contains two functionally important helices: an AH and a TMH.

### Ice2 family members typically have 10 transmembrane helices

Our identification of residues 245-262 as an AH not a TMH suggested further revision of Ice2p topology may be needed. We also noted that the C-terminus of the region between TMH7 and TMH8 (residues 379-400) forms another hydrophobic, conserved, helical region with biophysical properties of a TMH (Figure 1B; Supplementary Table 1), and intermediate predictions of forming a TMH in TMHMM2.0 (Supplementary Figure 2A). To investigate further, we applied other prediction tools. Phobius predicted 7 TMHs, one of which was residues 379-397 (not shown).^44^ TOPCONS predicted a consensus topology identical to TMHMM2.0. ^26^ However, three of the five prediction methods used by TOPCONS to generate a consensus predicted a TMH at residues 375-395, and two tools predicted a TMH at AH1, with low reliability reported for the whole region 240-400 (Supplementary Figure 2B).

**Figure 2.**
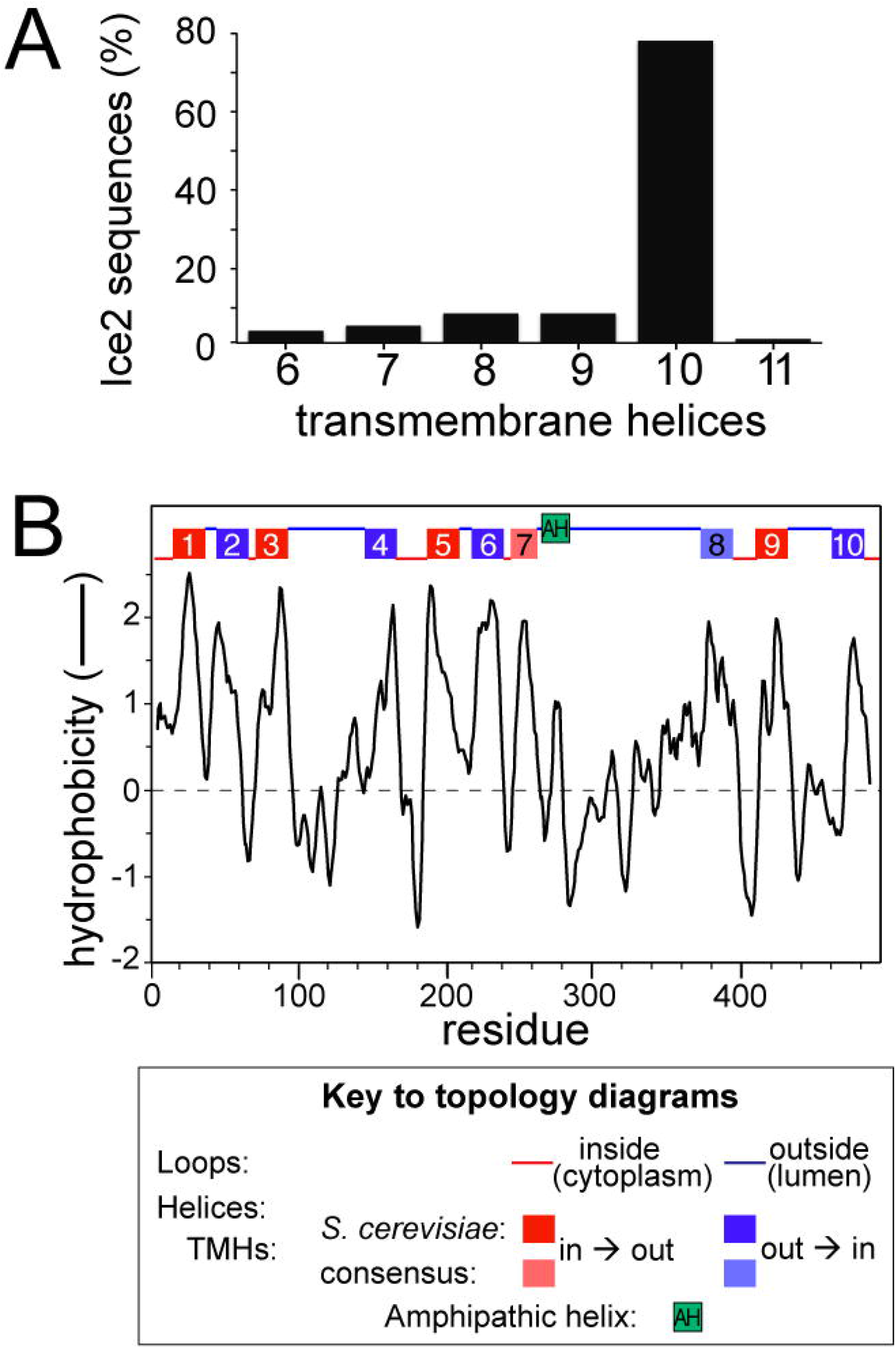
10 TMH topology from multiple sequence alignment across the Ice2 family. A. Number of predicted TMHs and B. Hydrophobicity among 95 aligned Ice2 sequences. In B, the average Kyte-Doolittle hydrophobicity was calculated using a sliding window of 11 residues. Topology of loops and TMH are indicated in the key.

As TMHMM and TOPCONS made inconsistent predictions, we switched to using MemBrain, a tool that shares some of their computational approaches (machine learning) and also has been developed in several ways, one being to include information from multiple homologous sequences.^28^ To promote a family-wide analysis, we excluded two non-conserved regions from Ice2p, reducing the sequence to 351 residues. The predicted topology was 10 TMHs: the same 8 as TMHMM and TOPCONS, plus the two we identified in the 240-415 region (Supplementary Figure 2C and Figure 1).

To further predict topology of Ice2p, we broadened the analysis beyond the *S. cerevisiae* protein, which is obviously variant for example in terms of insertions, to include proteins across the whole family (n=95). Given that Ice2 proteins form a single region of homology across their entire length (Supplementary Figure 1), we looked for a consensus topology across the protein family. 10 TMHs were predicted in 77% of fungal species (Figure 2A). The additional two TMHs not predicted by TMHMM/TOPCONS aligned with the hydrophobic regions we had already identified, as illustrated by plotting average hydropathy across the Ice2 multiple sequence alignment (Figure 2B). This strongly indicates that Ice2p in *S. cerevisiae* spans the ER membrane with 10 TMHs. One additional effect of this prediction is that it suggests an alternate location for the loop with its sole AH: the increased number of TMHs and predominant prediction of orientation places the AH in the ER lumen (Figure 2B).

### Profile-sequence searches detect homology between Ice2p and SERINCs

Although the Ice2 family is only present in fungi, we wondered if Ice2 has previously unrecognised distant homologues. All hits in a converged PSI-BLAST profile were already identified as Ice2 (not shown). We therefore switched to HMMER, a more sensitive profile-sequence search tool that uses hidden Markov models (HMMs) to build profiles.^45^ Iterative HMMER searches converged rapidly (3 iterations), with >99% of hits identified as Ice2 homologues and 0.5% of hits proteins without known domains in diverse non-fungal eukaryotes (Supplementary Table 2A). We then used HHblits, which is more sensitive and precise than HMMER.^46^ As with HMMER, the hit profile increased rapidly in iterations 1/2, but then grew slowly – approximately 5% per iteration for next 6 iterations, before expanding rapidly in iterations 9 to 11 (Supplementary Table 2C). The proteins added in iterations 3–8 were proteins of no known domain in widely dispersed eukaryotes, similar to the minority of proteins found by HMMER. A single hit in iteration 8 was documented as TMS1, from a large family ubiquitous in eukaryotes better known by a newer name: SERINCs, with five members in humans, and the sole yeast member called Tms1p.^20,47^ The rapid expansion in subsequent iterations 9+ resulted solely from SERINCs, which dominated numerically from iteration 11.

We next asked if the two budding yeast-specific inserts reduced effectiveness of profile-search tools. A profile seeded with Ice2 missing the two inserts (reduced to 351 aa) grew three times faster after iteration 2, becoming dominated by SERINCs from iteration 5 onwards (Supplementary Table 2D). Given the increased detectability of homologues using reduced Ice2, we then seeded HMMER with it, and found this search now behaved like HHblits, identifying SERINCs after a middle phase of slow growth of the set of proteins with no known domains (Supplementary Table 2B). In summary, Ice2p was identified as homologous to SERINC by both HHblits (TMHs 4-7, residues 140-260) and HMMER (TMHs 2-6, residues 40-230).

### Profile-profile searches confirm homology between Ice2p and SERINCs

To confirm the Ice2-SERINC homology, we initially carried out reverse searches by seeding profiles with SERINCs. However all of PSI-BLAST, HMMER and HHblits focussed entirely on SERINCs with no additional hits (not shown). This might arise from the numerical dominance of SERINC over the other sequences involved here.

We next made the searches more symmetrical, treating seed and target equivalently by using the profile-profile search tool HHsearch (implemented online as HHpred), which compares HMM-built profiles of both seeds and targets.^30,31^ Pairwise comparison of a bespoke multiple sequence alignment (MSA) that contains only SERINC with another that contains only Ice2 showed that the two MSAs aligned with estimated probability of a true positive match up to 98.6%, and expected value of alignment based on sequence alone to occur by random (E-value) down to 2×10^−9^ (Figure 3A, Supplementary File 1). In an open search of HMM databases made for every entry in PFAM or human or yeast proteomes, SERINCs and Ice2 identified each other as the sole strong hits apart from themselves, with all weaker hits showing a step change of decreasing strength length or both (Figure 3B). The region of Ice2 showing homology was residues 40-260 (TMHs 2–7). In addition, a different profile-profile tool, FFAS, produced the same links between the SERINC and Ice2 families, and were almost full-length (Supplementary Table 3).

**Figure 3.**
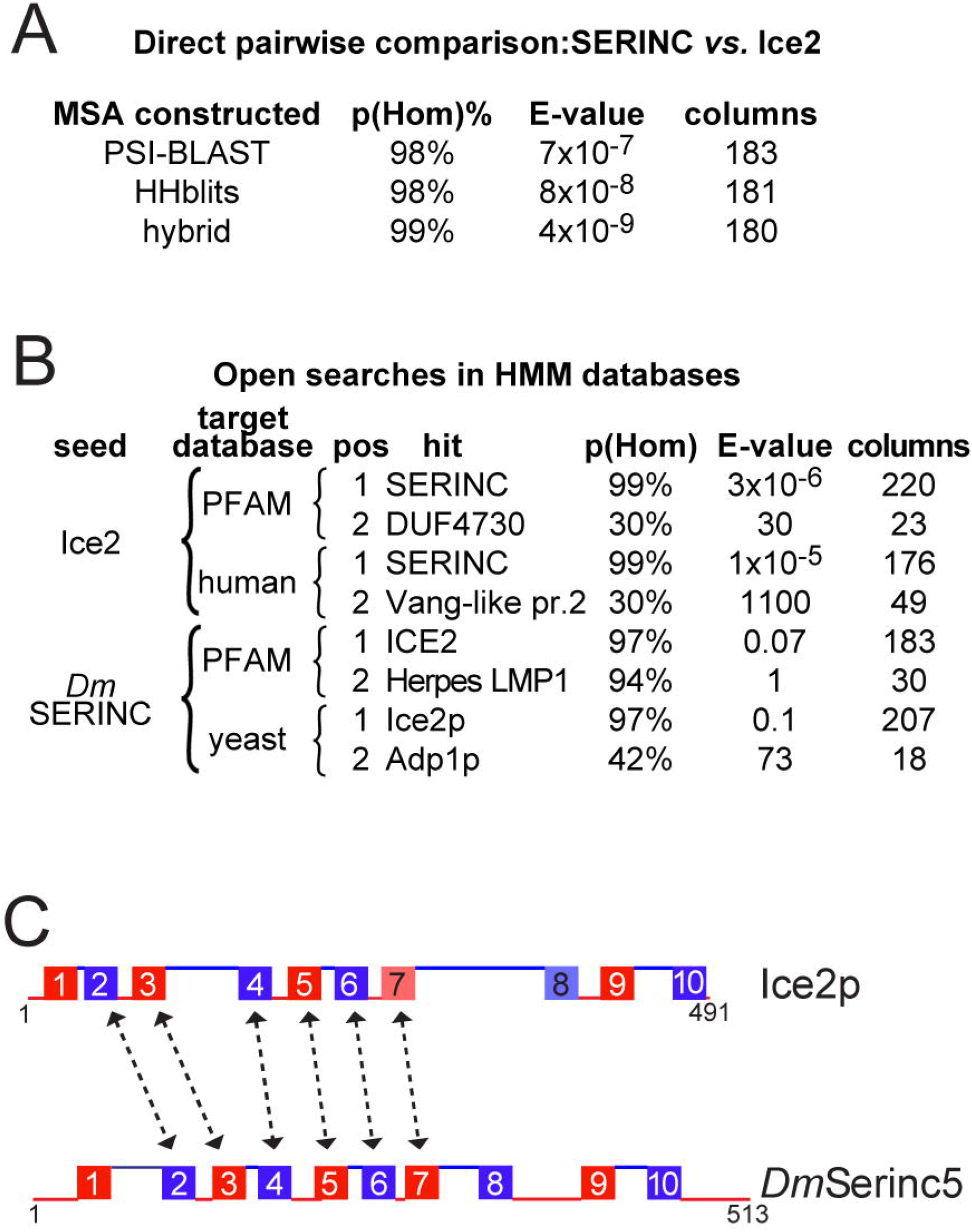
Profile-profile alignments of Ice2p with SERINCs. A. Probabilities reported by HHpred for SERINCs to be homologous to Ice2p (pHom, %), expected value for alignment of sequence of strength observed to occur by chance (E-value) and number of residues aligned (columns), as determined by HHpred for profile-profile alignments, with profiles made three ways: PSI-BLAST alone (nr70 and nr50 databases for Ice2p and SERINC respectively), HHblits alone (UniClust30 database, iteration before SERINCs entered the MSA), and hybrid as HHblits except the Ice2 profile was expanded from 61 to 1202 sequences by one round of BLAST into NCBI’s nr100 database, excluding SERINCs. B. Open search of all HMMs in databases made up of PFAM (18200 HMMs) and the human and yeast proteomes (109000 and 6002 HMMs respectively), showing top two non-self hits. C. Regions conserved between Ice2p (with topology from Figure 2) and SERINC5. Loops and TMHs indicated as in Figure 2B.

The Ice2 and SERINC homology is bolstered by the overall topological similarity between them, as SERINCs also have 10 TMHs (Supplementary Figure 3).^20^ The alignment is between TMHs 2-7 in register (Figure 3C). In addition, when we forced full-length alignment in HHpred (by reducing the threshold for “greediness”) additional sequence similarities were found, particularly in TMH 10 (Supplementary File 1). The hit in HHpred has six of the properties of a true positive, although it lacks a short linear motif.^48^ Overall, these results indicate that Ice2p is a variant of SERINC.

### An intermediate ICE2-like family identifies likely early evolutionary branching of Ice2 ancestor

To examine the evolutionary relationship between Ice2p and SERINCs, we studied the group of proteins with no known domains identified above by HMMER and HHblits. PSI-BLAST searches seeded with any one of these produced a profile in early iterations that contained the same group of 42 proteins mostly in animals (invertebrates, including chordates, for example *Branchiostoma*, and some insects), and also in dispersed unikonts, including one fungal sequence and several amoebae. This ability of all sequences to identify each other indicates that they form a previously unrecognised protein family.

Submitting PSI-BLAST profiles of this family for further iterations led to inclusion first of Ice2 family members (from iteration 2) then SERINCs (from iteration 4). HMMER profiles made the same links more rapidly than PSI-BLAST (SERINCs from iteration 3). Using a group of sequences enriched in the new family (Supplementary Table 4), we visualised its relationships with SERINCs and Ice2 both as a phylogeny tree and as a cluster map. Both visualisations showed that the new family is most closely related to Ice2. The new family and Ice2 share a common branch from SERINC (Figure 4A), and the new family is more similar to SERINCs than the Ice2 family is (Figure 4B). This identifies the new family as an intermediate between SERINC and Ice2. The most parsimonious explanation for the observed relationships is that an ancestor of the intermediate family diverged from ubiquitous SERINCs at an early stage of unikont evolution, and subsequently Ice2 developed within this family and became fixed in a fungal ancestor. Importantly, the ability to link Ice2 and SERINCs by PSI-BLAST that is seeded with any member of the new family shows that Ice2-SERINC homology is not an artefact of using HMM profiles.

**Figure 4.**
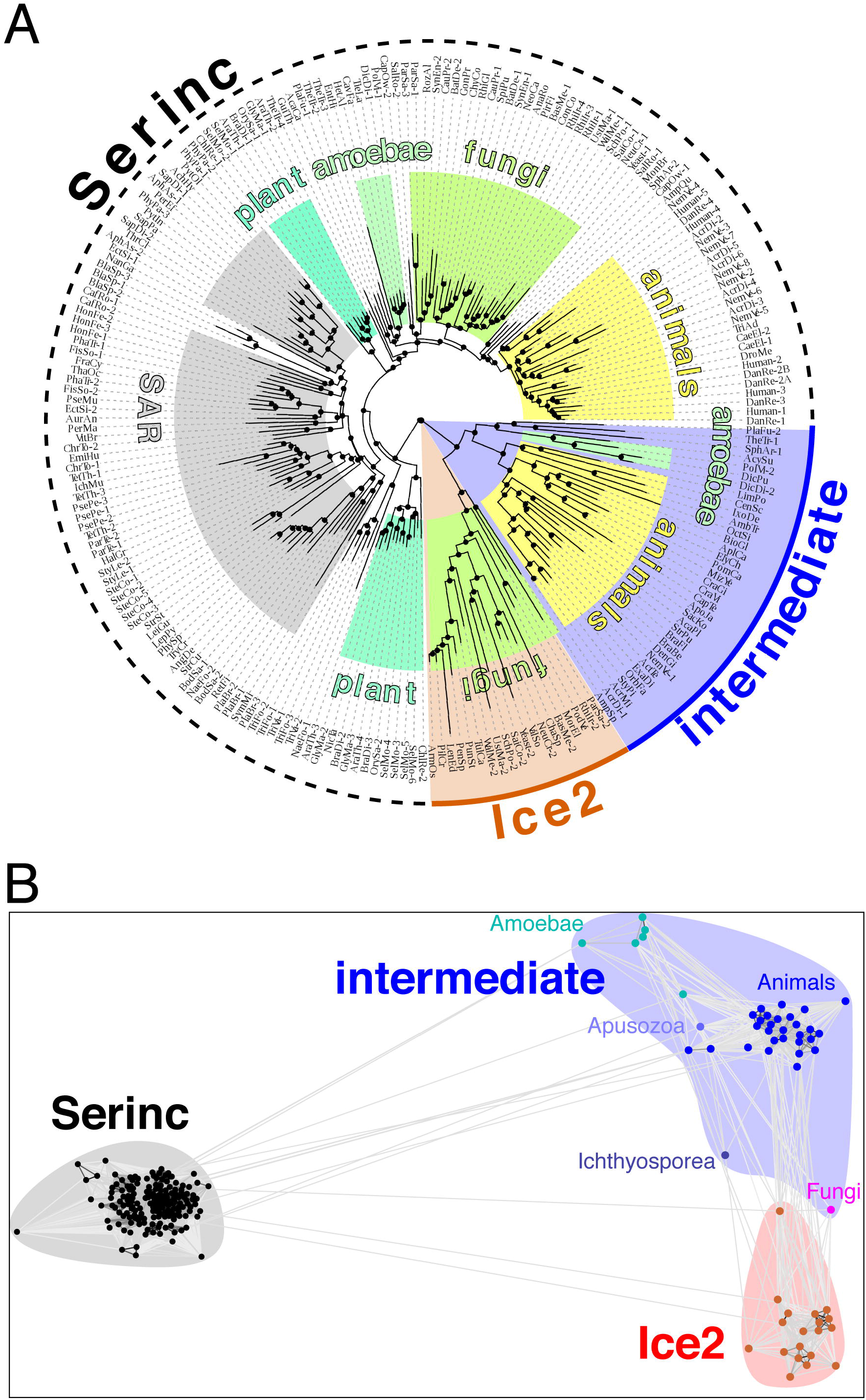
Relationships between SERINC, Ice2 and a previously unknown intermediate family. Phylogeny tree of three proteins families: SERINC, Ice2, and the new intermediate family. Cluster map of same proteins. Both diagrams show relationships for 223 sequences consisting of 171 diverse SERINCs, 19 Ice2s and 33 sequences not previously identified with any domain (see Supplementary Table 4). Protein families are coloured: SERINC: black, Ice2: red, intermediate family: blue. Additional colours in (A) indicate major taxonomic groupings in the tree. Additional colours in (B) indicate 5 clades within the intermediate family.

### Folding by inter-residue distance suggests Ice2-SERINC homology covers at least TMH1-9

We next used tools that fold proteins *ab initio* from pairwise co-evolution of contacting residues to determine if homology extended to encompass beyond TMHs 2-7. First, we compared the contact maps of the two proteins made by trRosetta. A contact map for *D. melanogaster* SERINC showed 18 strong pairwise inter-TMH contacts (Supplementary Figure 4A). These are compatible with the arrangement of TMHs observed by single particle cryo-EM structure of *D.m.* SERINC at a resolution of 3.3Å (PDB: 6SP2).^22^ This identified two subdomains: TMH1/2/3/9 (subdomain A) and TMH5/6/7/10 (subdomain B), with TMHs 4 and 8 crossing between subdomains (see Figure 5A).^22^ The contact map for Ice2 shares 13 of the 18 strong inter-TMH contacts (Supplementary Figure 4B). The exceptions involve TMHs 4, 8 and 10. For TMH10, in SERINC it contacts TMHs 4/6/7 on one face of subdomain B as well as a portion of TMH9, while in Ice2 it only shares strong contacts with TMHs7/9 and has an additional contact with TMH2, compatible with being in subdomain A. While SERINC shows three distinct contacts for portions of TMHs 4 and 8, for Ice2 these contacts do not stand out above background. The relative lack of clarity for Ice2 may result from the Ice2 MSA generated by trRosetta having only 30% of the number of sequences (n=239) as were included for SERINC (n=760). We therefore created a contact map for Ice2 with a much larger MSA, using RaptorX which accepts bespoke MSAs (Supplementary Figure 4C).^49^ This supported the existence of the 13 strong interactions found by trRosetta to be shared with SERINC, and confirmed that TMH10 differs in Ice2p from SERINC, as it does not interact with TMH4, now interacting with TMH1 (not TMH2).

**Figure 5.**
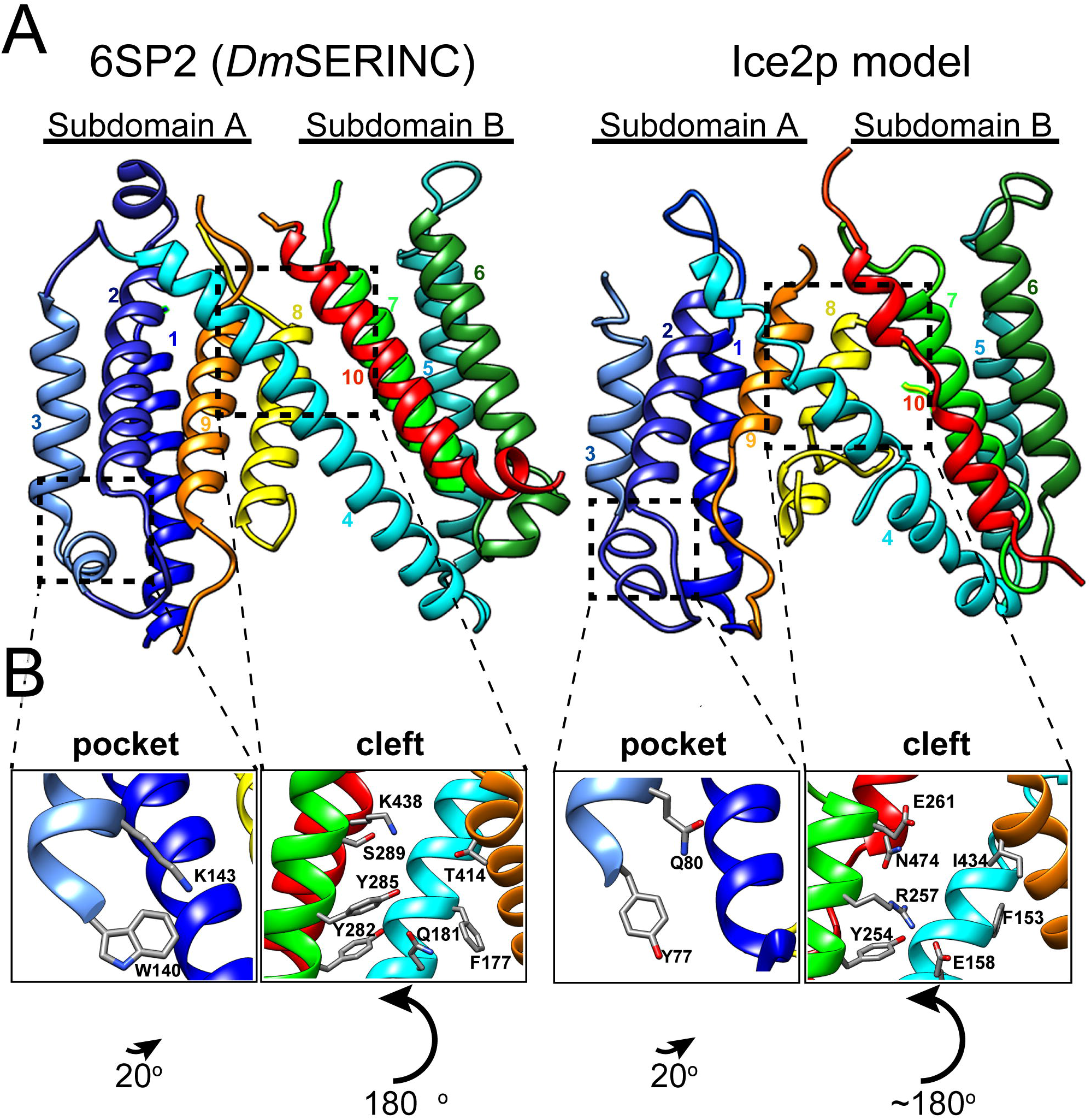
Structural model of Ice2p. A. The Cryo-EM structure of *Dm*SERINC (6SP2) compared to a model of Ice2p. Transmembrane helices are numbered and coloured in a spectrum from blue (N-terminus) to red (C-terminus). The Ice2p model was generated by Modeller, based on an HHpred alignment that maximally extends the homology between Ice2p and 6SP2 (see also Supplementary File 1). B. Zoomed in view of two structural elements in *Dm*SERINC and corresponding regions in the Ice2p model: (i) pocket on the cytoplasmic face of subdomain A, showing TMH1/3, with 2 conserved residues; (ii) cleft at the back, lumenal side where Subdomains A and B meet, made up of TMH4/7/9/10 and TMH8 (removed), with a hydrophobic phenylalanine plug at the base and lined by hydrophilic residues.

Overall, the finding that SERINC and Ice2 share most of the strong TMH interactions estimated by pairwise evolution supports a much wider homology between Ice2 and SERINC than identified by profile alignment tools.

### 3D-modelling of Ice2 identifies key conserved residues likely to have functional roles in SERINCs

Other template-based tools (Phyre2, Galaxy and Swiss-Model) all found that Ice2 makes a significant top hit to SERINC, with homology for the region TMHs 2-7 (not shown).^34–36^ We also used the threading tool I-TASSER.^50^ This made a full-length model of Ice2 (with loops deleted) based on the SERINC structure, with RMSD 1.7Å across 311 residues and a TM-score of 0.53, where >0.5 signifies that the structures come from the same superfamily (Figure 5A).

The Ice2 model shared two structural features with SERINC that have been proposed to be functionally important: a cleft with a hydrophilic lining in the lumenal side of the protein between the two subdomains; and a pocket on the cytoplasmic face of subdomain A. Among the 16 key residues identified in these two sites (8 lining the cleft and 8 in the pocket), 9 are well-aligned with Ice2 in the profile-profile alignment by HHpred (Supplementary File 1). 9/16 represents a significant enrichment, compared to 73 well-aligned residues across all 331 columns (Chi^2^_n−1_ p=7×10^−4^), showing that the alignment focussed on residues with likely functional roles. These key residues can be visualised in a model, including F154 in Ice2p, plugging the base of the cleft and polar side chains of E158/Y254 lining the cleft, aligning with F177 and Q181/ Y282 respectively in *Dm*SERINC (Figure 5B). Likewise for the pocket, Y77 and Q80 in Ice2p align with W140 and K143 in SERINC (Figure 5B).

## DISCUSSION

### Ice2p is a member of the SERINC Superfamily

Our aim was to use bioinformatics to learn more about the function for Ice2p, an integral ER membrane protein of unknown function in yeast. First, we revised the predicted topology of Ice2p to 10 TMHs. This required comparisons to be made across the fungal Ice2 family, most likely because the budding yeast protein is an outlier (for example, with inserts), which put topology predictions with this single sequence on the borderline of reliability (not shown). Next, we found that Ice2p is a full-length homologue of SERINCs, another family of membrane proteins of unknown function at the molecular level, these being distributed across all eukaryotes, often with multiple members per species (for example, five in humans). The homology between individual yeast Ice2p and SERINC sequences is distant (12% identity overall, Supplementary File 1), but using the whole Ice2 family to build up a profile creates homology that is so strong that it can be established using PSI-BLAST, so long as the search is enriched with the newly identified intermediate family (Figure 4).

Homology can also be established with any seed if HMM-based profile-sequence tools are used, though HMMER searches with Ice2 make the link if the *S. cerevisiae*-only inserts are removed from Ice2p (Supplementary Table 2). Starting searches with SERINC, PSI-BLAST fails because of low variance and large size (not shown), and HMMER almost makes the link, so that at convergence (18th iteration, 4081 hits), one of the 63 non-significant hits (1> E-value >0.01) was an Ice2 (not shown). 12% identity is typical of the homology generally required for profile-profile tools.^30,51^ HHpred identifies the link very clearly, reporting 99% probability of homology across six TMHs, and a highly significant E-value based on sequence alone (4×10^−9^, Figure 3). Among the seven criteria that distinguish between false and true positives in HHpred, the only one that Ice2 lacks is a conserved motif.^48^ This may be because the key functional elements are all in the intra-membrane TMHs, so residues that function together are separated by 3 (or 2) intervening residues.

Other tools corroborated the main result, including the independent approach of pairwise co-evolution. This showed that nine of the helices in Ice2p are arranged similarly to those in SERINC (Supplementary Figure 4). The contacts made by TMH10 in Ice2 appeared different from SERINC, suggesting a possible different position. However, the N-terminus TMH10 still preserved contact with TMH9 and the C-terminus with TMH6/7 suggesting that TMH10 runs along the same course (Figure 5). This fits with TMH10 showing sequence conservation at the same for higher levels than any other TMH (Supplementary File 1). Finally, threading by I-TASSER extended homology to full-length. Together, this evidence makes a strong prediction that adds the Ice2 family to a newly appreciated SERINC superfamily.

The origin of Ice2 can be seen to some extent from our discovery of a family intermediate between Ice2 and SERINC. Members are mostly in invertebrates, but one is in a fungus, and others are in amoebae (*i.e.* outside opisthokonts) (Figure 4). This suggests that this family and Ice2 branched from SERINC in a unikont common ancestor, and that Ice2 expanded within fungi under tight sequence constraints.

While it is possible that Ice2p has a similar activity to SERINC, the degree of functional relatedness is not yet known. While 9 of the 16 key residues were identified as well conserved, there are non-aligned residues in both the cleft and pocket, in which one of the key residues is completely missing: TMH9 in Ice2 is missing a helical turn (4 residues) where SERINCS have a conserved intra-TMH histidine (Supplementary File 1). Thus it is possible that function have been repurposed across the SERINC superfamily. While the Ice2 model cannot determine its function, it can guide experiments by demonstrating key residues and surfaces.

### Ice2p is predicted to interact with specific lipids

Structural and mechanistic studies of SERINC allow useful predictions about Ice2p based on its membership of the SERINC superfamily.^22,52^ Initially, SERINCs were proposed to modulate membrane lipid composition.^20^ However, lipidomic studies have failed to detect the impacts of SERINC1 or SERINC5 on membrane composition of phosphatidylserine or sphingolipids.^53,54^ This suggests that SERINCs exert their antiviral actions through other mechanisms. The structure of *Drosophila* SERINC shows a cleft in the lumenal leaflet that is accessible to solvent molecules in molecular dynamics simulations, with a nearby hydrophobic groove between TMHs 6/7/8 occupied by an acyl chain in the structure.^22^ In addition, specific lipids (cholesterol, phosphatidylserine and sulphatide) stabilised *Drosophila* SERINC against breakdown during heating, suggesting that these lipids might specifically bind the protein.

The structural homology includes not only the hydrophilic cleft and the pocket, but also the hydrophobic groove (not shown). This suggests that like SERINC, Ice2p has an intramembrane lipid binding site, though the lipid involved is unknown. This might explain the repeated linking of Ice2p to phospholipid synthesis,^14,52^ and may be related to Ice2’s direct binding of the Pah1 phosphatase regulatory subunit Spo7.^16^

### Ice2p is unlikely to be a conventional tether

Ice2p has been proposed to tethering the ER to lipid droplets through four AHs in a long cytoplasmic TMH6/7 loop that targeted lipid droplets when expressed on its own.^14^ The findings here predict Ice2 has identical topology to SERINC, which places the loop (probably with one single AH) inside the lumen of the ER between TMHs 7/8. This implies that the loop might access lipid droplets *in vivo* only if multiple TMHs change topology. Reversible topology changes have been reported for bacterial permeases and transporters, and they have been predicted in eukaryotes,^55^ with a few examples reported.^56,57^ This makes it is worth considering whether Ice2p, and also SERINC, might have reversible topology.

Along with multiple ER-related functions attributed to Ice2p,^8,10,11,13,14^ it is required for normal formation of cortical ER in multiple different genetic backgrounds, including wild-type (only buds affected),^5^ cells with excess cortical ER because of unregulated phospholipid synthesis (*opi1*Δ),^16^ and cells lacking cortical ER from other lesions.^6,7^ Our results call into question the idea that Ice2p forms a physical bridge (“tether”) to the plasma membrane.^7^. Even if the AH flips into the cytoplasm, it does not have the polybasic nature of helices that can bind the plasma membrane.^58,59^ An alternative means of bridging would be a plasma membrane protein binding partner for Ice2, but none was found in a proximity screen.^16^ A more parsimonious hypothesis is that Ice2p modulates a fundamental aspect of ER function and/or regulates other ER proteins, for example the Nem1/Spo7 phosphatase complex, and hence phospholipid biosynthesis and overall ER size, which is most variable in the cortical region.^16^ This may parallel the ability of SERINCs in viral envelopes to regulate other membrane proteins, for example indirectly modulating the conformation and clustering of HIV Env.^52^

### Yeast as a future model for SERINC research

Although more is known at the molecular level about SERINCs than Ice2p ^22,52^, identifying the link to Ice2p may aid research into SERINCs. *S. cerevisiae* is typical of fungi that express an Ice2 family member in that it also expresses a SERINC: Tms1p. Little is known about this protein, except that it localises to the degradative vacuole (equivalent of lysosome) ^60^, and that it can bind a limited range of lipids ^61^. Targeting of multiple subcellular compartments by a pair of yeast SERINC superfamily proteins resembles the wide intracellular distribution of human SERINCs to plasma membrane ^21^, ER ^20^, and pre-autophagic vesicles ^62^. To date, attempts to produce a SERINC-null organism have been limited to a high-throughput study of *C. elegans*, where double RNAi of its two recently duplicated SERINCs produced no synthetic negative interaction ^63^. Future yeast studies with individual and double *ice2/tms1* deletion mutants may lead to improved understanding of the whole superfamily.

## Supporting information

All supplementary Material (4 Figs, 4 Tables and 1 Supplementary File)

## Acknowledgements

The work was funded by grant BB/P003818/1 from the Biotechnology and Biological Sciences Research Council (BBSRC), UK

## Contributions

G.O.A. and T.P.L. were both responsible for Conceptualisation, Formal Analysis, Writing (Original Drafted Review/Editing) and Visualisation.

## Supplemental Information

4 Supplemental Figures, 4 Supplemental Tables and one Supplemental File.

## Supplementary Figure Legends

**Supplementary Figure 1: Phylogenetic tree and sequence alignment in the Ice2 family.**

A. Phylogenetic relationships between Ice2p and homologues from 17 other fungi dispersed across evolution. B. Alignment of the 18 Ice2 sequences by Clustal Omega. Colouring of conserved residues is by CLUSTALX.

**Supplementary Figure 2: Topology of Ice2p reported by different tools.**

**A.** Topology analysis of *S. cerevisiae* Ice2p by TMHMM2.0 (Krogh et al., 2001). For key, see Figure 2B. **B.** TOPCONS report on Ice2p. TOP: Membrane topologies of Ice2p as predicted with reports of consensus (top) and five constituent tools, with key to describe loops and TMHs. The position of the sole strongly predicted AH is indicated as in part A. BOTTOM: Reliability of overall consensus, only reporting values above 0.9. **C**. MemBrain 3.1 report on Ice2p. Here, two non-conserved loops (246-266 and 379-400, indicated by yellow blocks in part B, the second of which overlaps with the AH) were excluded. Loops and TMHs are indicated as in B. In addition to 10 TMHs, the prediction includes one “half TMH” of 7 residues before TMH4.^64^

**Supplementary Figure 3: Alignment of SERINCs from key organisms across evolution**

Alignment of 17 SERINC sequences from human (×5), *D. melanogaster* (×1), *C. elegans* (×2), *S. cerevisiae* (×1), *S. pombe* (×1), *D. discoideum* (×1), *C. reinhardtii* (×2), and *A. thaliana* (×4). Colouring is according to the CLUSTALX scheme. Positions of TMHs 1-10, the N-terminal leader, and long loop after TMH8 are indicated. Number of residues removed in long unique inserts are indicated in square brackets.

**Supplementary Figure 4: Contact maps for Ice2 and SERINC**

A and B. Maps made by trRosetta for SERINC (using yeast Tms1p) and full length Ice2 respectively. C. RaptorX. For A and B the map reports the predicted distance of Cb atoms between 20Å and 4Å with coloured scale from white<->yellow<->red<->black. For C, the map reports probability of contact, defined as distance between backbone carbons ≤8Å, with darker grey indicating higher probability. In all parts, the position of the TMHs and other structural elements are indicated along the axes. Also indicated are the major contacts between TMHs or segments of TMHs (N/C termini and M= middle). Contacts are shown in black writing in a grey box, except if only found in Tms1p (blue), or only found in Ice2p (red). Annotations are placed in the bottom/left half of each diagram, leaving the symmetric top/right half left unannotated. A and B are additionally annotated to identify boundaries of secondary structural elements (dashed lines), and two other elements: location of TMHs on the main diagonal (dotted boxes); possible regions of interaction by non-TMH regions (light yellow fill).

