## Supplementary material for "Fungal Ice2p is in the same superfamily as SERINCs, restriction factors for HIV and other viruses": All supplementary Material (4 Figs, 4 Tables and 1 Supplementary File)

A) Supplementary Figures 1-4: pages (1-6)
B) Supplementary Tables 1-4: 6 pages (7-12)
C) Supplementary File 1: 1 page (13)

#### A) Legends for Supplementary Figures

Supplementary Figure Legends

**Supplementary Figure 1: Phylogenetic tree and sequence alignment in the Ice2 family.**

A. Phylogenetic relationships between Ice2p and homologues from 17 other fungi dispersed across evolution. B. Alignment of the 18 Ice2 sequences by Clustal Omega. Colouring of conserved residues is by CLUSTALX. *S. cerevisiae* Ice2p is highlighted in yellow.

**Supplementary Figure 2: Topology of Ice2p reported by different tools.**

**A.** Topology analysis of *S. cerevisiae* Ice2p by TMHMM2.0 (Krogh et al., 2001). For key, see Figure 2B. **B.** TOPCONS report on Ice2p. TOP: Membrane topologies of Ice2p as predicted with reports of consensus (top) and five constituent tools, with key to describe loops and TMHs. The position of the sole strongly predicted AH is indicated as in part A. BOTTOM: Reliability of overall consensus, only reporting values above 0.9. **C**. MemBrain 3.1 report on Ice2p. Here, two non-conserved loops (246-266 and 379-400, indicated by yellow blocks in part B, the second of which overlaps with the AH) were excluded. Loops and TMHs are indicated as in B. In addition to 10 TMHs, the prediction includes one “half TMH” of 7 residues before TMH4.[64]

**Supplementary Figure 3: Alignment of SERINCs from key organisms across evolution**

Alignment of 17 SERINC sequences from human (x5), *D. melanogaster* (x1), *C. elegans* (x2), *S. cerevisiae* (x1), *S. pombe* (x1), *D. discoideum* (x1), *C. reinhardtii* (x2), and *A. thaliana* (x4). Colouring is according to the CLUSTALX scheme. Positions of TMHs 1-10, the N-terminal leader, and long loop after TMH8 are indicated. Number of residues removed in long unique inserts are indicated in square brackets.

**Supplementary Figure 4: Contact maps for Ice2 and SERINC**

A and B. Maps made by trRosetta for SERINC (using yeast Tms1p) and full length Ice2 respectively. C. RaptorX. For A and B the map reports the predicted distance of Cb atoms between 20Å and 4Å with coloured scale from white<->yellow<->red<->black. For C, the map reports probability of contact, defined as distance between backbone carbons ≤8Å, with darker grey indicating higher probability. In all parts, the position of the TMHs and other structural elements are indicated along the axes. Also indicated are the major contacts between TMHs or segments of TMHs (N/C termini and M= middle). Contacts are shown in black writing in a grey box, except if only found in Tms1p (blue), or only found in Ice2p (red). Annotations are placed in the bottom/left half of each diagram, leaving the symmetric top/right half left unannotated. A and B are additionally annotated to identify boundaries of secondary structural elements (dashed lines), and two other elements: location of TMHs on the main diagonal (dotted boxes); possible regions of interaction by non-TMH regions (light yellow fill).

### Supplementary Figure 1

A

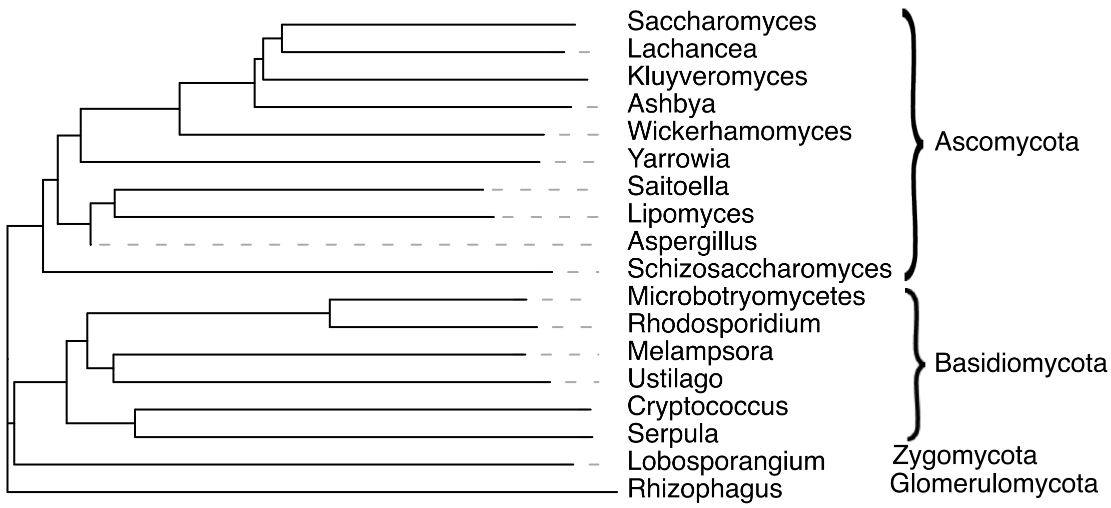


B

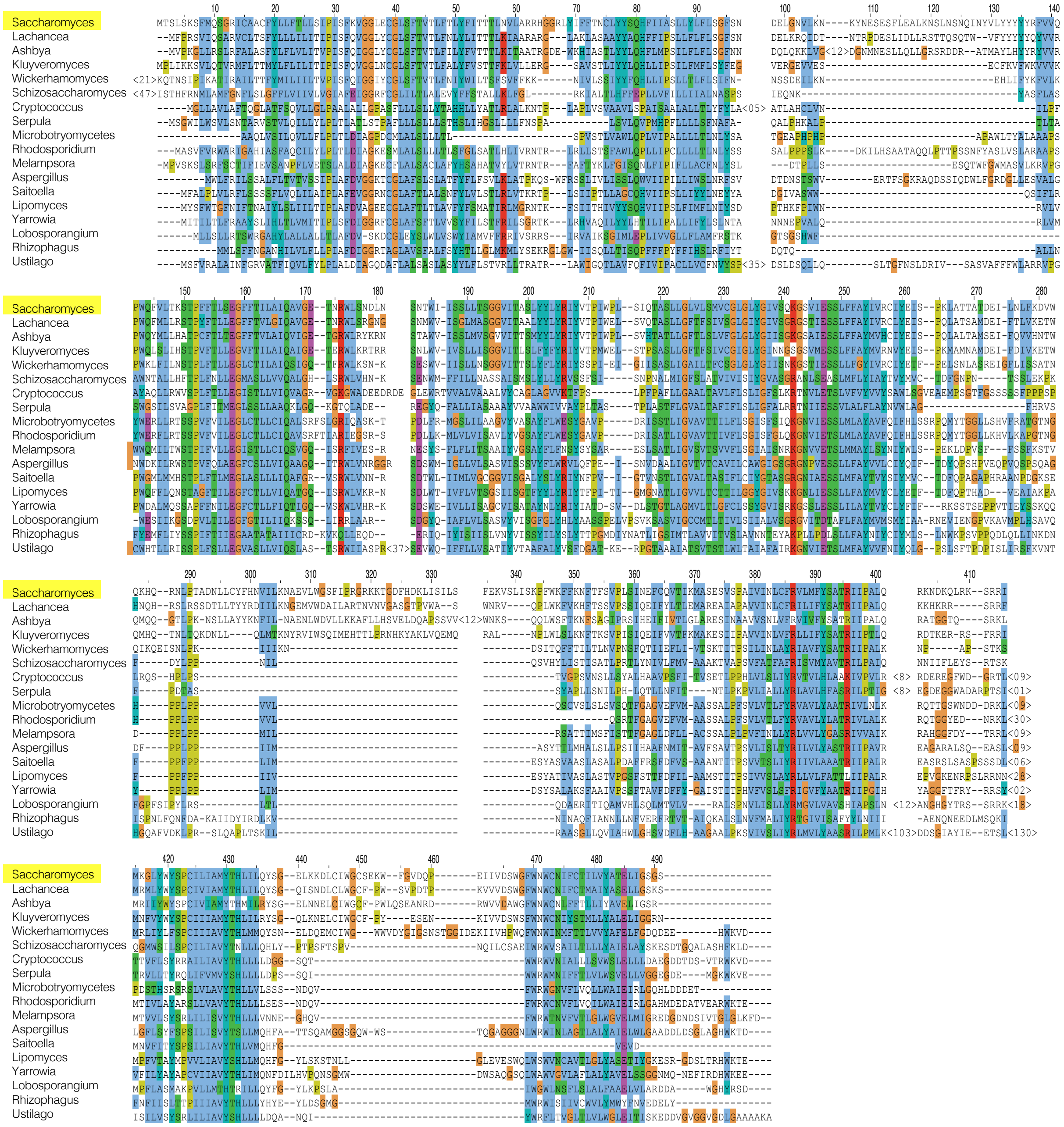

### Supplementary Figure 2

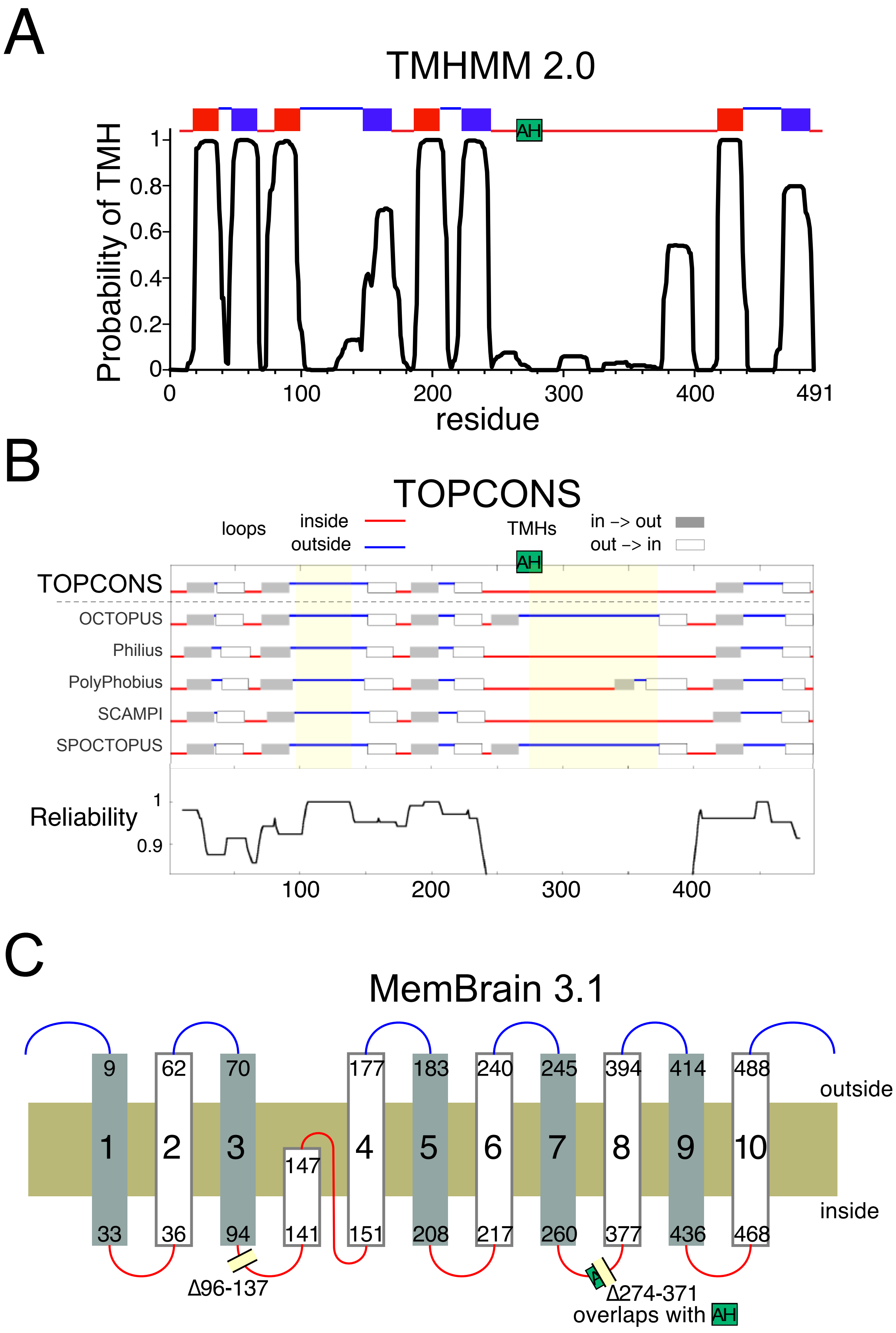

### Supplementary Figure 3

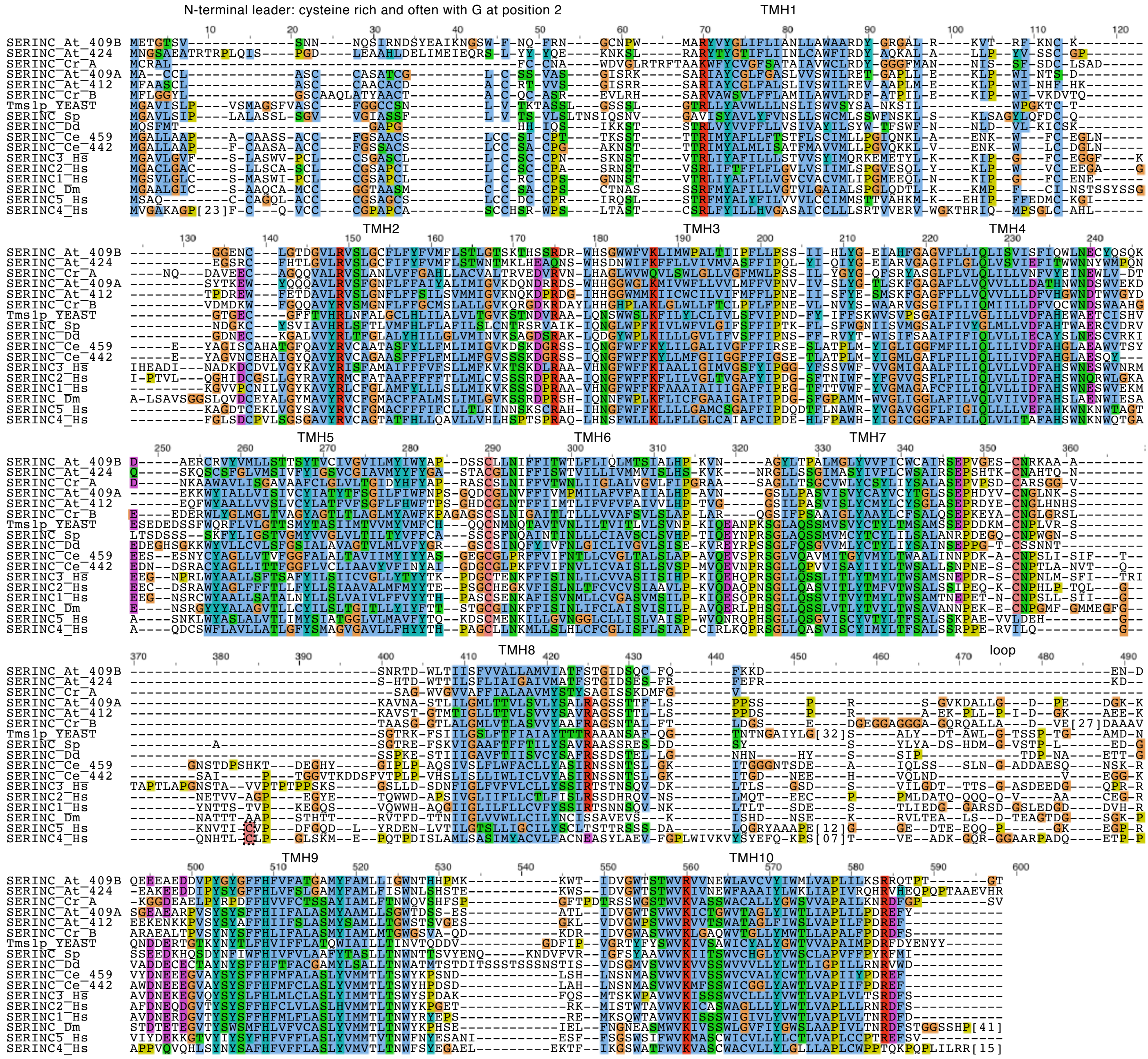

### Supplementary Figure 4

#### A. Tms1p - trRosetta

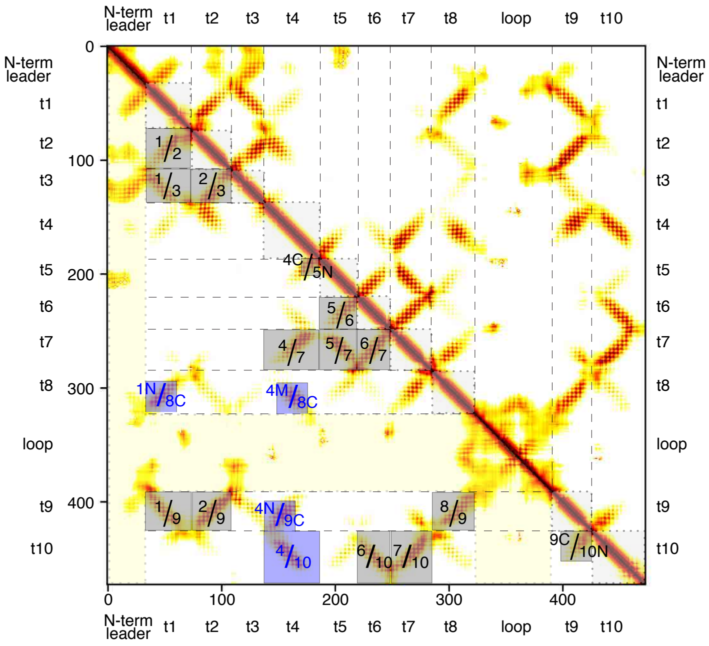


#### B. Ice2p - trRosetta

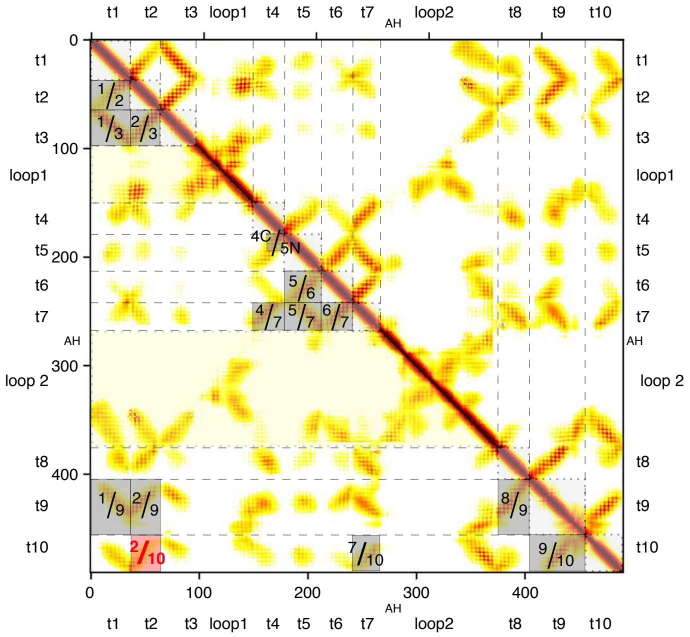


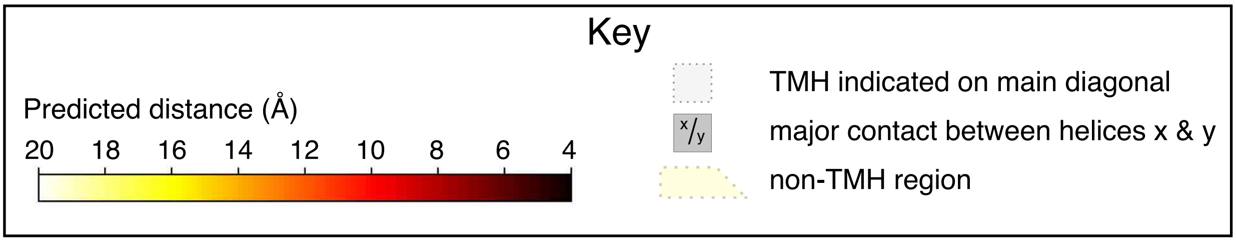


#### C. Ice2 (loops removed) - RaptorX

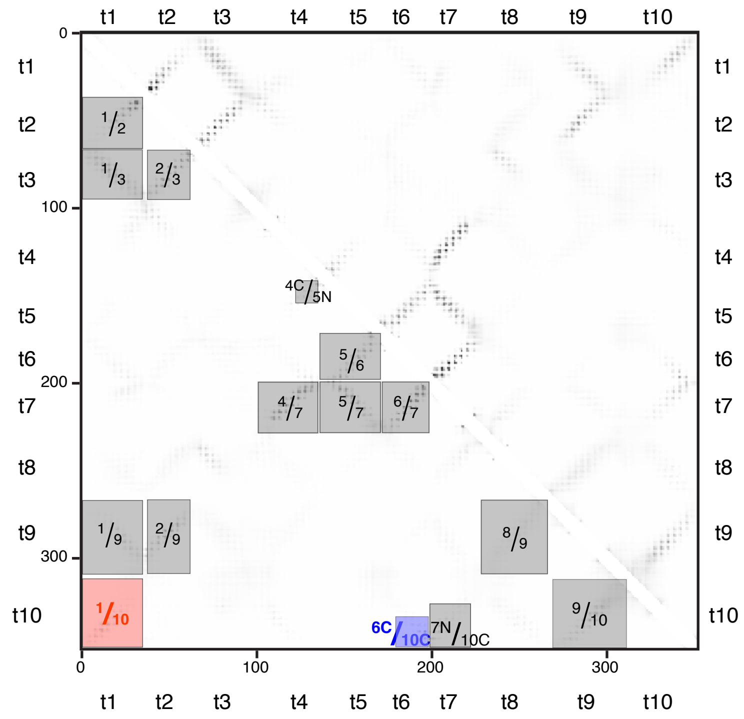

### Supplementary Tables 1-4

#### Supplementary Table 1

Properties of predicted TMHs and AHs in Ice2p

| Ice2p | residues | Hydro-phobicity | Hydrophobic moment | % polar residues |
|---|---|---|---|---|
| *Previously predicted transmembrane helices (TMH) in Ice2p* | | | | |
| **TMH1** | 14-31 | 1.19 | 0.32 | 17% |
| **TMH2** | 37-54 | 1 | 0.1 | 39% |
| **TMH3** | 68-85 | 1.02 | 0.21 | 28% |
| **TMH4** | 152-169 | 0.91 | 0.20 | 33% |
| **TMH5** | 185-202 | 0.87 | 0.21 | 44% |
| **TMH6** | 222-239 | 1.01 | 0.15 | 33% |
| **TMH7** | 418-435 | 1.19 | 0.1 | 17% |
| **TMH8** | 467-484 | 1.14 | 0.24 | 28% |
| *Previously predicted amphipathic helices (AH)* | | | | |
| **AH1** | 245-262 | 0.91 | 0.29 | 28% |
| **AH2** | 266-283 | 0.53 | 0.50 | 50% |
| **AH3** | 293-310 | 0.64 | 0.39 | 39% |
| **AH4** | 333-350 | 0.66 | 0.40 | 44% |
| *Additional conserved region* | | | | |
| **after AH4 (see Figure 1C)** | 383-400 | 0.93 | 0.09 | 22% |
| *Helical properties of amphipathic helices with known targeting preferences* | | | | |
| **Lipid droplet (VPS13A)** | 2991-3008 | 0.40 | 0.42 | 56% |
| **Plasma membrane (Ist2p)** | 936-946* | 0.18 | 0.60 | 36% |

The helical parameters for each highlighted structural feature in Ice2p (all 18 residues) were generated by the Heliquest tool (https://heliquest.ipmc.cnrs.fr/; Gautier et al., 2008), and compared to AHs that target lipid droplets (VPS13A) and plasma membrane (Ist2p) (Ercan et al., 2009; Kumar et al., 2018). Values are coloured: yellow if characteristic of TMHs (hydrophobicity ≥0.8, hydrophobic moment ≤0.3, polar residues ≤40%), blue if characteristic of AHs (hydrophobicity ≤0.6, hydrophobic moment ≥0.4, polar residues ≥50%) and grey if intermediate between these values.

Supplementary Table 2

HMM-based profile searches for Ice2 either full length (A/C) or with 96-137 / 274-371 omitted (B/D) using HMMER (A/B) and HHblits (C/D).

A. Ice2p (HMMER)

| iter-ation | Total hits | New Ice2 |
|---|---|---|
| 1 | 514 | 514 |
| 2 | 605 | 91 |
| 3 | 610 | 5 |
| 4 | 610 | 0 |

B. Ice2 missing 2 inserts (HMMER)

| iter-ation | Total hits | New Ice2 | New SERINC | New Others |
|---|---|---|---|---|
| 1 | 518 | 518 | 0 | 0 |
| 2 | 608 | 88 | 0 | 2 |
| 3 | 614 | 0 | 0 | 6 |
| 4 | 622 | 0 | 0 | 8 |
| 5 | 627 | 0 | 1 | 4 |
| 6 | 641 | 0 | 14 | 0 |
| 7 | 2680 | 0 | 2039 | 0 |

C. Ice2p (HHblits)

| iter-ation | Total hits | New Ice2 | New SERINC | New others |
|---|---|---|---|---|
| 1 | 32 | 32 | 0 | 0 |
| 2 | 51 | 19 | 0 | 1 |
| 3 | 53 | 1 | 0 | 1 |
| 4 | 55 | 0 | 0 | 2 |
| 5 | 59 | 0 | 0 | 4 |
| 6 | 64 | 0 | 0 | 5 |
| 7 | 68 | 0 | 0 | 4 |
| 8 | 70 | 0 | 1 | 1 |
| 9 | 87 | 0 | 17 | 0 |
| 10 | 151 | 0 | 64 | 0 |
| 11 | 328 | 0 | 177 | 0 |

D. Ice2 missing 2 inserts (HHblits)

| iter-ation | Total hits | New Ice2 | New SERINC | New Others |
|---|---|---|---|---|
| 1 | 33 | 33 | 0 | 0 |
| 2 | 50 | 16 | 0 | 1 |
| 3 | 61 | 1 | 0 | 10 |
| 4 | 70 | 0 | 2 | 7 |
| 5 | 138 | 0 | 68 | 0 |

For each iteration, hits are classified from the UNIPROT record as Ice2, Serinc or other. The class that grows the most is highlighted in yellow. Only Ice2p (full-length) in HMMER (part A) onverged in the number of iterations shown.

Supplementary Table 3

FFAS03 was seeded with Ice2p and human SERINC5, reporting the top three hits in both PFAM and yeast databases, with scores below -9.5 being significant, and thos e withhigher scores in grey.{Jaroszewski, 2005 #3019}

| Seed | Database | Hit | Score | Template | %id | First | Last |
|---|---|---|---|---|---|---|---|
| Ice2p | PFAM | 1 | -105 | PF08426; ICE2 | 28 | 1 | 417 |
| | | 2 | -17 | PF03348; Serinc | 10 | 19 | 443 |
| | | 3 | -7.04 | PF15256; SPATIAL | 16 | 156 | 198 |
| | Yeast | 1 | -140 | Ice2p | 100 | 1 | 464 |
| | | 2 | -15.8 | Tms1p | 11 | 39 | 459 |
| | | 3 | -9.59 | Pex30p | 8 | 24 | 351 |
| Serinc | PFAM | 1 | -103 | PF03348; Serinc | 24 | 1 | 450 |
| | | 2 | -15.2 | PF08426; ICE2 | 10 | 24 | 416 |
| | | 3 | -8.81 | PF07415; herpes LMP2 | 15 | 136 | 459 |
| | Yeast | 1 | -109 | Tms1p | 23 | 1 | 471 |
| | | 2 | -14.6 | Ice2p | 9 | 9 | 461 |
| | | 3 | -10.4 | Cks1p | 10 | 9 | 150 |

Supplementary Table 4

**223 sequences used to demonstrate relationships in the SERINC superfamily.**

Sequences were obtained from three sources as described in the Methods: (1) 184 proteins from a converged PSI-BLAST seeded with a member of the new family. The number of hits was kept low by restricting hits with 29 taxonomic terms (in yellow below); (2) 34 proteins with no known domains were amalgamated from the hits of the 2nd iteration of 2 PSIBLAST searches seeded with members of the new family; (3) 5 sequences curated by hand from incomplete hits in groups (1) and (2).

Short names used in the tree are 3 and 2 letters from the two species names, except human and *S. cerevisiae* ("yeast"). For species with more than one sequence, numbering is as indicated.

**(1) Converged PSI-BLAST (9 iterations) with *T. trahens* XP_013756618.1 restricted to 29 taxonomic groups (highlighted yellow), plus 4 fungal groups: Chytridiomycota, Cryptomycota, Mucoromycota, Zoopagomycota, then edited down to 184 sequences.**

Animals:- *Acropora digitifera*: XP_015753109.1 (AcrDi-1); XP_015779446.1 (AcrDi-2); XP_015761684.1 (AcrDi-3); XP_015771525.1 (AcrDi-4); XP_015769349.1 (AcrDi-5); XP_015747507.1 (AcrDi-6); *Amphimedon queenslandica*: XP_003382411.1; *Caenorhabditis elegans*: NP_506611.1 (CaeEl-1); NP_741561.1 (CaeEl-2); *Danio rerio*: NP_956021.1 (DanRe-1); NP_001038647.1 (DanRe-2A); XP_009290780.1 (DanRe-2B); XP_003199457.1 (DanRe-3); XP_698642.3 (DanRe-4); *Drosophila melanogaster*: NP_001261947.1; *Homo sapiens*: AAQ88795.1 (Human-1); NP_001185967.1 (Human-2); AAB48858.1 (Human-3); NP_001244960.1 (Human-4); XP_011541606.1 (Human-5); *Nematostella vectensis*: XP_001631941.1 (NemVe-1); XP_032228872.1 (NemVe-2); XP_001633416.1 (NemVe-3); XP_001625482.2 (NemVe-4); XP_032236962.1 (NemVe-5); EDO43486.1 (NemVe-6); XP_032238402.1 (NemVe-7); XP_001631880.2 (NemVe-8); *Trichoplax adhaerens*: XP_002114508.1.

Choanoflagellata:- *Monosiga brevicollis*: XP_001745243.1; *Salpingoeca rosetta*: XP_004994555.1 (SalRo-1); XP_004997867.1 (SalRo-2).

Filasterea:- *Capsaspora owczarzaki*: XP_011270789.1 (CapOw-1); XP_004346817.1 (CapOw-2).

Ichthyosporea:- *Sphaeroforma arctica*: XP_014155466.1 (SphAr-1); XP_014156139.1 (SphAr-2).

Fungi:- *Anaeromyces robustus*: ORX85449.1; *Basidiobolus meristosporus*: ORY01015.1 (BasMe-1); ORX86092.1 (BasMe-2); *Batrachochytrium dendrobatidis*: XP_006680018.1 (BatDe-1); XP_006681464.1 (BatDe-2); *Caulochytrium protostelioides*: RKP03832.1 (CauPr-1); RKO97024.1 (CauPr-2); *Chytriomyces confervae*: TPX75829.1; *Conidiobolus coronatus*: KXN65508.1; *Gonapodya prolifera*: KXS20326.1; *Neocallimastix californiae*: ORY77155.1; *Neurospora crassa*: XP_962656.1 (NeuCr-1); XP_960816.1 (NeuCr-2); *Paramicrosporidium saccamoebae*: PJF19746.1; PJF19195.1 (ParSa-1); PJF18006.1 (ParSa-3); *Piromyces finnis*: ORX50272.1; *Rhizoclosmatium globosum*: ORY36319.1; *Rhizophagus irregularis*: XP_025167307.1 (RhiIr-1); XP_025183532.1 (RhiIr-2); XP_025185281.1 (RhiIr-3); XP_025168662.1 (RhiIr-4); *Rozella allomycis*: RKP20934.1; *Saccharomyces cerevisiae*: NP_010390.3 (Yeast-1); NP_012176.1 (Yeast-2); *Saitoella complicata*: XP_019025563.1 (SaiCo-1); XP_019024036.1 (SaiCo-2); *Schizosaccharomyces pombe*: *sp*|Q9HDY3.1|*YK17_SCHPO* (SchPo-1); *sp*|O13909.1|YDW1_*SCHPO* (SchPo-2); *Spizellomyces punctatus*: XP_016611289.1; *Synchytrium endobioticum*: TPX43735.1 (SynEn-1); TPX43303.1 (SynEn-2); *Ustilago maydis*: XP_011388355.1 (UstMa-1); XP_011386607.1 (UstMa-2); *Wallemia mellicola*: XP_006956946.1 (WalMe-1); XP_006958532.1 (WalMe-2).

Apusozoa:- *Thecamonas trahens*: XP_013756618.1 (TheTr-1); XP_013754690.1 (TheTr-2); XP_013756847.1 (TheTr-3); XP_013753078.1 (TheTr-4).

Amoebozoa:- *Acanthamoeba castellanii*: XP_004336528.1; *Acytostelium subglobosum*: XP_012753995.1; *Cavenderia fasciculata*: XP_004366977.1; *Dictyostelium discoideum*: XP_640818.1 (DicDi-1); XP_642737.1 (DicDi-2); *Entamoeba histolytica*: GAT97894.1; *Heterostelium album*: XP_020429608.1; *Planoprotostelium fungivorum*: PRP77133.1 (PlaFu-1); PRP86022.1 (PlaFu-2); *Polysphondylium violaceum*: KAF2073863.1 (PolVi-1); KAF2073390.1 (PolVi-2); *Tieghemostelium lacteum*: KYQ89646.1.

Metamonada:- *Blastocystis sp.*: XP_014526547.1 (BlaSp-1); OAO13453.1 (BlaSp-2); XP_014529570.1 (BlaSp-3); *Trichomonas vaginalis*: XP_001329577.1 (TriVa-1); XP_001304501.1 (TriVa-2); *Tritrichomonas foetus*: OHT06334.1 (TriFo-1); OHS95406.1 (TriFo-2); OHS99326.1 (TriFo-3).

Discoba:- *Angomonas deanei*: EPY32215.1; *Bodo saltans*: CUG92822.1 (BodSa-1); CUI12368.1 (BodSa-2); *Guillardia theta*: XP_005841735.1; *Leishmania guyanensis*: CCM16931.1; *Leptomonas pyrrhocoris*: XP_015663863.1; *Phytomonas sp.*: CCW63503.1; *Strigomonas culicis*: EPY32228.1; *Trypanosoma cruzi*: XP_821245.1.

Plants:- *Arabidopsis thaliana*: NP_187268.2 (AraTh-1); CAA0210062.1 (AraTh-2); NP_001324781.1 (AraTh-3); NP_001328593.1 (AraTh-4); *Brachypodium distachyon*: XP_010237604.1 (BraDi-1); KQJ83457.1 (BraDi-2); XP_010230536.1 (BraDi-3); *Chlamydomonas reinhardtii*: PNW79375.1 (ChlRe-1); XP_001703677.1 (ChlRe-2); *Glycine max*: XP_003552131.1 (GlyMa-1); XP_006605737.1 (GlyMa-2); XP_003530635.1 (GlyMa-3); *Oryza sativa*: EAY87831.1 (OrySa-1); EEC70050.1 (OrySa-2); *Selaginella moellendorffii*: XP_024518071.1 (SelMo-1); XP_024528794.1 (SelMo-2); EFJ32526.1 (SelMo-3); XP_024543165.1 (SelMo-4); EFJ22865.1 (SelMo-5); XP_024526619.1 (SelMo-6).

Haptista:- *Chrysochromulina tobinii*: KOO29417.1 (ChrTo-1); KOO29447.1 (ChrTo-2); *Reticulomyxa filosa*: ETO12023.1.

SAR:- *Achlya hypogyna*: OQR99138.1; *Aphanomyces astaci*: XP_009827937.1 (AphAs-1); XP_009832039.1 (AphAs-2); *Aureococcus anophagefferens*: XP_009040848.1; *Cafeteria roenbergensis*: KAA0153407.1 (CafRo-1); KAA0145593.1 (CafRo-2); *Ectocarpus siliculosus*: CBN74600.1 (EctSi-1); CBJ27137.1 (EctSi-2); *Emiliania huxleyi*: XP_005765279.1; *Fistulifera solaris*: GAX24251.1 (FisSo-1); GAX16780.1 (FisSo-2); *Fragilariopsis cylindrus*: OEU13331.1; *Halteria grandinella*: TNV78744.1; *Hondaea fermentalgiana*: GBG27723.1 (HonFe-1); GBG24993.1 (HonFe-2); GBG30373.1 (HonFe-3); *Ichthyophthirius multifiliis*: XP_004030116.1; *Naegleria fowleri*: KAF0983897.1 (NaeFo-1); KAF0974805.1 (NaeFo-2); *Nannochloropsis gaditana*: EWM24934.1; *Paramecium tetraurelia*: XP_001433369.1 (ParTe-1); XP_001441224.1 (ParTe-2); *Perkinsus marinus*: XP_002772482.1; *Peronospora effusa*: RQM09469.1; *Phaeodactylum tricornutum*: XP_002179611.1 (PhaTr-1); XP_002180794.1 (PhaTr-2); *Phytophthora palmivora*: ETK85404.1 (PhyPa-1); POM80097.1 (PhyPa-2); ETI51004.1 (PhyPa-3); *Plasmodiophora brassicae*: CEO97761.1 (PlaBr-1); CEO97071.1 (PlaBr-2); SPQ97588.1 (PlaBr-3); *Pseudo-nitzschia multistriata*: VEU43570.1; *Pseudocohnilembus persalinus*: KRX09925.1 (PsePe-1); KRX07118.1 (PsePe-2); KRX10953.1 (PsePe-3); *Pythium insidiosum*: GAY00482.1; *Pythium oligandrum*: TMW62873.1; *Saprolegnia diclina*: XP_008615013.1 (SapDi-1); XP_008608891.1 (SapDi-2); *Saprolegnia parasitica*: XP_012203268.1; *Stentor coeruleus*: OMJ92313.1 (SteCo-1); OMJ69345.1 (SteCo-2); OMJ90402.1 (SteCo-3); OMJ73821.1 (SteCo-4); OMJ89300.1 (SteCo-5); *Streblomastix strix*: KAA6404213.1; *Stylonychia lemnae*: CDW84436.1 (StyLe-1); CDW86304.1 (StyLe-2); *Symbiodinium microadriaticum*: OLQ04137.1; *Tetrahymena thermophila*: XP_001009070.1 (TetTh-1); XP_001032827.2 (TetTh-2); XP_001012741.1 (TetTh-3); *Thalassiosira oceanica*: EJK73866.1; *Thraustotheca clavata*: OQS02657.1; *Vitrella brassicaformis*: CEM38767.1.

**(2) 34 sequences from 2nd iteration of PSI-BLAST searches seeded with *T. trahens* XP_013756618.1 and *N. vectensis* XP_001631941.1.**

Animals:- *Acanthaster planci*: XP_022104596.1; *Acropora millepora*: XP_029184522.1; *Actinia tenebrosa*: XP_031573879.1; *Aplysia californica*: XP_005094950.1; *Apostichopus japonicus*: PIK53311.1; *Biomphalaria glabrata*: XP_013096175.1; *Branchiostoma belcheri*: XP_019622340.1; *Capitella teleta*: ELT98757.1; *Crassostrea gigas*: XP_011414590.1; *Crassostrea virginica*: XP_022335780.1; *Dendronephthya gigantea*: XP_028396389.1; *Elysia chlorotica*: RUS85734.1; *Exaiptasia pallida*: XP_020894608.1; *Limulus polyphemus*: XP_022243097.1; *Mizuhopecten yessoensis*: XP_021373130.1; *Octopus vulgaris*: XP_029642825.1; *Orbicella faveolata*: XP_020622096.1; *Pomacea canaliculata*: XP_025091768.1; *Saccoglossus kowalevskii*: XP_006822995.1; *Strongylocentrotus purpuratus*: XP_030831010.1; *Stylophora pistillata*: XP_022780854.1.

Fungi:- *Amphiamblys sp.*: OIR59099.1; *Armillaria ostoyae*: SJL05432.1; *Chaetothyriales sp.*: RMZ75936.1; *Lentinula edodes*: GAW06119.1; *Mortierella elongata*: OAQ30346.1; *Podila verticillata*: KFH62670.1; *Peniophora sp.*: VDB84951.1; *Piloderma croceum*: KIM85142.1; *Punctularia strigosozonata*: XP_007378719.1; *Tulasnella calospora*: KIO18403.1; *Valsa sordida*: ROV98548.1.

**(3) Further 5 sequences added by hand**

Animals:- *Amblyomma triste*: A0A023G3K0; *Branchiostoma floridae*: XP_035691277.1; *Centruroides sculpturatus*: XP_023230516.1 & XP_023230491.1; *Ixodes ricinus*: A0A131XSX6.

Plants:- *Nicotiana Tabacum*: XP_016507244.1.

### Supplementary File 1

#### Alignment of Ice2p with D. melanogaster SERINC by HHpred

```
                                TMH1                                                               core start        TMH2
S 2°str     CchhHHHHHHHHHHHHHHHHHH----HHHHHHH----------------HH----hhc----c ChHHHHHHHHHHHHHHH
Ice2        MTSLSKSFMQSGRICAACFYLL----FTLLSIP----------------IS----FKV----G GLECGLSFTVTLFTLYF 52
S Cons.     ~~~~~~~~~~~~~~~~~~~~~~----~ill~lP----------------L~----f~~----~ G~~~~L~~Sl~L~~~~~
            ++|.+-.+..++.++.+.+...    ..+--+|                +.    ++.    . |+++|||+||++++||+
T Cons.     ~~r~~y~~~~~~~~~~~~~~~~~~~~~~~~l~~~~~~~~~~~~~~~~~~~~~~~~~~~~~~C~~~~ g~~~vyR~~~~~~~ff~
DmSERINC    SSRFMYAFILLVGTVLGAIALSPGLQDTLKKMPFCINSTSSYSSGALSAVSGGSLQVDCEYAL GYMAVYRVCFGMACFFA 116
T 2°str     HHHHHHHHHHHHHHHHHHHHcCcchHHHHhcCCcccCCCCccCcchhhhccCCCccCcHHHHH HHHHHHHHHHHHHHHHH

                                             TMH3               Δ42                ‡ TMH4
S 2°str     HHHHHHHHcccCccchHHHHHHHHHHHHHHHHHHHHHHHHHHhh-c*hhccHHHHHHHHHhcChHHHHHHHHHHHHHHHHHHHHHH
Ice2        ITTTLNVLARRHGGRLYIFFTNCLYYSQHFIIASLLYLFLS-G*FFVVQPWQFVLTKSTPFFTLSEGFFTILAIQAVGET 172
S Cons.     i~~tl~l~~~~~~~~~~~~~~~~~~l~~lq~~~~~~lL~~~l~-~*~~~~~~w~~~L~~s~P~F~llEG~~sLlvIq~~gq~
            +++++++.+++.++ +|+.+||++|.+|+++++++++++|+ | +. |..+|.++.++++.+|+++|.+.++|++|.|||
T Cons.     ~~~~~~~~~~~~~~~-~r~~~h~g~w~~k~~~~~~~~~~~f~ip~~~~~~~~~~~~~~~~~~fi~iq~i~~idf~~~~~~-
DmSERINC    LMSLIMLGVKSSRD-PRSHIQNNFWPLKFLICFGAAIGAIFIPDGSFGPAMMWVGLIGGLAFILVQLVIIVDFAHSLAE- 194
T 2°str     HHHHHHcCCCCCCC-HHHHHHhccHHHHHHHHHHHHHHHHHHhCCCccchhHHHHHHHHHHHHHHHHHHHHHHHHHHHHHH-

                                       TMH5                                   TMH6
S 2°str     HHHHHhcc-CCchHHHHHHHHHHHHHHHHHHHHHH---HHHHcCCCCCCCHHHHHHHHHHHHHHHHHHHHHHHHc------CC
Ice2        NRWLSNDL-NSNTWIISSLLTSGGVITASLYYL---YRIYVTPIWPLSIQTASLLGLVLSMVCGLGLYGIVS------QK 242
S Cons.     ~r~li~~~-~s~~~~i~~Li~Sa~~y~~s~~~l---y~iy~~p~~~~s~~~a~LlG~~lt~~v~L~~~g~~~-------~r
            +|+||+ +++.|+..++..+.++|++|++++ |   .+|+++   ++|..|.++|+++++++++++++++++|       |+
T Cons.     --w~~~~~~~~~~w~~~l~~~~~~~~~~~~~~~~~~~~~~~~~--~~C~~n~~~i~~~~~l~~~~~~~~s~~p~~~~~~~~
DmSERINC    --NWIESAENSRGYYYALAGVTLLCYILSLTGITLLYIYFTTS--TGCGINKFFISINLIFCLAISVISILPAVQERLPH 270
T 2°str     --HHHHHhhhcCcccHHHHHHHHHHHHHHHHHHHHHHHHHHHHhhcc--cCCchHHHHHHHHHHHHHHHHHHHHHHHcHHHhcCCCC

                         TMH7      core end                                Δ98                   TMH8
S 2°str     CChhHHHHHHHHHHHHHHHH ----------HCchhhc------C-Cc---hh-**-HhcCCHHHHHHHHHHHH-HHHHHHHHH
Ice2        GSVIESSLFFAYIVRCIYE ----------ISPKLAT------T-AT---DE-**-SESVSPAIVINLCFRV-LMFYSATRI 396
S Cons.     g~iiEssll~~yiv~~iy~ ----------~~~~~~~------~-~~---~~-**-~~~l~~~vlisL~yRl-~vl~~asrI
            ++++++|++++|++|++|+           ||+++.+      + .+    |.    +.+.++..++++++.+ +++|+++|.
T Cons.     ~gll~s~~v~~y~~yl~~s al~~~p~~~cn~~~~~~~~~~~~~~~~~~~~~~~~~~~~~~~~~~i~~~~~~~~~~~~y~~~~~
DmSERINC    SGLLQSSLVTLYTVYLTWS AVANNPEKECNPGMFGMMEGFGNATTTAAPSTHTTRVTFDTTNIIGLVVWLLCILYNCISS 350
T 2°str     CcchHHHHHHHHHHHHHHH HHHcCCccccChhHHHhhcCCCCCccccCCCCCCCCCCCcHHHHHHHHHHHHHHHHHHHHHhc

                                                                         ‡          TMH9
S 2°str     HH---HHHhcCc-----------c-cccc-------cccchhHHHHHHHH----HHHHHHHHHHHHHHhhcccccccccC
Ice2        IP---ALQRKND-----------K-QLRK-------SRRIMKGLYWYSPC----ILIAMYTHLILQYSGELKKDLCIWGC 450
S Cons.     ip---~i~~~~~-----------~-d~~~-------~~~~~~~~~~ysp~----ilI~VYT~Lll~~~~~~~~~~~~~~~
            ++   +++..+|           + |||+       +||++++.|+|+.|.    ++++||++|++|||+.++.+  .+..
T Cons.     ~~~~~~~~~~~~~~~~~~~~~~~~~~~~~~~~~~~~~~~~~~~e~~~~~ysy~~fh~~~~las~y~~m~ltnW~~~~~~--~~~~
DmSERINC    AVEVSKISHDNSEKRVLTEALSDTEAGTDGSGKPSTDTETEGVTYSWSMFHLVFVCASLYVMMTLTNWYKPHSE--IELF 428
T 2°str     ccchhccCCCcccccchhhhhccCCCCCCCCCCCCCCccccCCCcCcHHHHHHHHHHHHHHHHHHHhcccCCchh--hhcc

                                      TMH10
S 2°str     cccccCCCCCccccCCchHHHHHHHHHHHHHHHHHHHHccC
Ice2        SEKWFGVDQPEIIVDSWGFWNWCNIFCTILVYATELIGSG 490 (491)
S Cons.     ~~~~~~~~~~~~~~~~~~~~WRWini~~tl~lY~~eL~~~~
            +++|              .++|+|++++|+|.+||+|+|++|.
T Cons.     ~~~~--------------~~~wvk~~s~w~~~~ly~wtl~ap~
DmSERINC    NGNE--------------ASMWVKIVSSWLGVFIYGWSLAAPI 457 (513)
T 2°str     CCCh--------------HHHHHHHHHHHHHHHHHHHHHHHhhH
```

Statistics: p(Hom): 98.6%; E-value: 4x $10^{-9}$; aligned cols: 180 core region (331 expanded); identities: 12%.

Multiple sequence alignments from 3 iterations of HHblits, with the Ice2 MSA being explanded by 1 round of BLAST against nr100, were compared pairwise with settings to align beyond the core region of homology. The 7 lines for the full alignment show:

Lines 2/6: sequences: Seed (S) = Ice2p 1-491; Target (T) *Dm*SERINC 29-458 (of 513), coloured by residue type: negative, positive, hydrophobic (incl proline), hydrophilic, others

Lines 1/7: predicted 2° structure: H=helix with numbering for Ice2p, E=strand, C=loop; CAPS=strong prediction

Lines 3/5: consensus sequences

Line 4: alignment at each residue: | strong; + good; . intermediate; - poor.

TMH1-10 are indicated along with core region (180 columns) that generated that calculated E-value.

Also indicated: key residues in SERINC (and matching residues in Ice2) lining the hydrophilic cleft (yellow) and the pocket on the cytosolic face of subdomain A (pink); and gaps in aligned TMHs (highlighed in blue). Inserts removed from Ice2 are identified by * (96-137) and by ** (274-371).
